# The commercial antimicrobial triclosan induces high levels of antibiotic tolerance *in vitro* and reduces antibiotic efficacy up to 100-fold *in vivo*

**DOI:** 10.1101/090829

**Authors:** Corey Westfall, Ana Lidia Flores-Mireles, John Isaac Robinson, Aaron J.L. Lynch, Scott Hultgren, Jeffrey P. Henderson, Petra Anne Levin

**Affiliations:** Department of Biology, Washington University in St. Louis, St. Louis, MO 63130, USA; Department of Molecular Microbiology and Microbial Pathogenesis, Washington University School of Medicine, St. Louis, MO 63110, USA; Division of Infectious Diseases, John T. Milliken Department of Internal Medicine, Washington University School of Medicine, St. Louis, MO 63110, USA; Department of Biological Sciences, University of Notre Dame, Notre Dame, Indiana, USA

**Author notes:** These two authors contributed equally to this study (Corey Westfall, Ana Lidia Flores-Mireles).

## Abstract

The antimicrobial triclosan is used in a wide range of consumer products ranging from toothpaste, cleansers, socks, and baby toys. A bacteriostatic inhibitor of fatty acid synthesis, triclosan is extremely stable and accumulates in the environment. Approximately 75% of adults in the US have detectable levels of the compound in their urine, with a sizeable fraction of individuals (>10%) having urine concentrations equal to or greater than the minimal inhibitory concentration for *Escherichia coli* and methicillin-resistant *Staphylococcus aureus* (MRSA). Previous work has identified connections between defects in fatty acid synthesis and accumulation of the alarmone guanosine tetraphosphate (ppGpp), which has been repeatedly associated with antibiotic tolerance and persistence. Based on these data, we hypothesized that triclosan exposure may inadvertently drive bacteria into a state in which they are able to tolerate normally lethal concentrations of antibiotics. Here we report that clinically relevant concentrations of triclosan increased *E. coli* and MRSA tolerance to bactericidal antibiotics as much as 10,000 fold *in vitro* and reduced antibiotic efficacy up to 100-fold in a mouse urinary tract infection model. Genetic analysis indicated that triclosan-mediated antibiotic tolerance requires ppGpp synthesis, but is independent of growth. These data highlight an unexpected and certainly unintended consequence of adding high concentrations of antimicrobials in consumer products, supporting an urgent need to reevaluate the costs and benefits of the prophylactic use of triclosan and other bacteriostatic compounds.

**Importance:** Added as a prophylactic to a wide range of consumer products, the fatty acid synthesis inhibitor triclosan accumulates to high levels in humans and the environment. Based on links between defects in fatty acid synthesis and accumulation of the alarmone ppGpp, we hypothesized that triclosan would render cells tolerant to bactericidal compounds due to ppGpp-mediated inhibition of biosynthetic capacity. Our data indicate that clinically relevant concentrations of triclosan induces higher tolerance of *E. coli* and methicillin resistant *S. aureus* (MRSA) to a panel of bactericidal antibiotics up to 10,000-fold. In a urinary tract infection model, mice exposed to triclosan exhibited bacterial loads ~100-fold higher in the bladder than control animals following ciprofloxacin challenge. These findings highlight an unexpected consequence of antimicrobials in consumer products and support an urgent need to reevaluate the costs and benefits of the prophylactic use of triclosan and other bacteriostatic compounds.

## Introduction

The prophylactic use of antibiotics in consumer goods, ranging from animal feed to personal care products, is widely believed to be a major contributor to the epidemic increase in antibiotic-resistant pathogens (1–3). Prominent among these prophylactics are triclosan and triclocarban, polychlorinated aromatic antimicrobials targeting fatty acid synthesis. Triclosan in particular is found in a wide variety of consumer products, including: toothpaste, cleansers, socks, and baby toys (1). Although the US Food and Drug Administration effectively banned the use of triclosan in household soap in 2017, as of this writing Canada and Australia, among other countries, have not elected to take similar action.

An inhibitor of enoyl-acyl carrier protein reductase, (4), at low concentrations (200 ng/ml) triclosan is bacteriostatic, preventing cell growth but having little effect on viability over the short term. At high concentrations (>10 μg/ml), triclosan is bactericidal, most likely killing cells through disruption of plasma membrane integrity (4). Recent work from the Waters lab suggests that at these higher concentrations triclosan can serve as an adjuvant, acting synergistically with tobramycin and other drugs, increasing killing by ~100-fold in a *Pseudomonas aeruginosa* biofilm model for CF lung infections (5). Triclosan is typically used as an antimicrobial additive at these higher, bactericidal concentrations.

Because of its widespread use as a prophylactic, the high concentrations at which it is employed, and its inherent stability, triclosan accumulates to high levels in the environment (6, 7). Approximately 75% of adults in the US have detectable levels of the compound in their urine, and >10% have urine concentrations greater than or equal to the minimal inhibitory concentration for *Escherichia coli* (200 ng/mL) and methicillin-resistant *Staphylococcus aureus* (MRSA) (100 ng/mL) (8, 9).

While the inverse relationship between antibiotic use and antibiotic efficacy is largely attributable to the selection of heritable traits, non-heritable traits such as antibiotic tolerance and persistence are also likely to be involved (10). In contrast to genetically-resistant bacteria, which grow in the presence of an antibiotic, tolerant bacteria are able to survive antibiotic challenge for longer periods of time than their more sensitive counterparts (10). Persister cells are the small sub-set of an otherwise-sensitive population (~1 in 10^6^) that exhibit levels of tolerance sufficient to protect them from otherwise lethal concentrations of antimicrobial compounds (11). Increases in antibiotic tolerance and persistence are confounding factors in the treatment of chronic *P. aeruginosa* (12) and *S. aureus* (13) infections and are thought to contribute to the refractory nature of medically relevant biofilms (14). Reduced growth rate and metabolic activity is associated with increased antibiotic tolerance (10) and is a defining trait of persister cells.

Based on previous work identifying connections between defects in fatty acid synthesis and accumulation of the alarmone guanosine tetraphosphate (ppGpp) (15), and the possible link between ppGpp and antibiotic tolerance (16), we hypothesized that triclosan exposure may inadvertently drive bacteria into a metabolically depressed state in which they are able to tolerate normally lethal concentrations of antibiotics (17, 18). In particular, inhibiting fatty acid synthesis stimulates interaction between acyl carrier protein and the hydrolase domain of the bifunctional ppGpp synthase SpoT, resulting in accumulation of the alarmone and the concomitant inhibition of biosynthetic capacity (19).

Here we report that clinically-relevant bacteriostatic concentrations of triclosan increased *E. coli* and methicillin resistant *Staphylococcus aureus* (MRSA) tolerance to bactericidal antibiotics as much as 10,000-fold *in vitro* and reduced antibiotic efficacy ~100-fold in a mouse urinary tract infection model. Triclosan-mediated antibiotic tolerance is dependent on ppGpp synthesis: although triclosan inhibited the growth of both wild-type and ppGpp mutant cells, only the latter were highly susceptible to challenge with bactericidal compounds. In contrast, pretreatment with another bacteriostatic drug, spectinomycin, a translation inhibitor that does not impact ppGpp accumulation (20), induced high levels of antibiotic tolerance in both wild-type and ppGpp mutant cells. Together, these data highlight an unexpected and certainly unintended consequence of employing triclosan as a commercial antimicrobial, and support an urgent need to reevaluate the costs and benefits of the addition of triclosan, and potentially other bacteriostatic compounds, to consumer products.

## Results

### Triclosan pretreatment results in high levels of tolerance to bactericidal antibiotics *in vitro*

To assess if physiologically relevant levels of triclosan are sufficient to promote tolerance to bactericidal antibiotics, we examined the relative sensitivity of *E. coli* (MG1655) and *S. aureus* (FPR3757 an USA-300 MRSA strain) cultured in minimal inhibitory concentrations (MIC) of triclosan to a panel of bactericidal antibiotics. Triclosan MICs for *E. coli* and MRSA were 200 ng/mL and 100 ng/mL, respectively under our growth conditions; similar to the triclosan concentration found in the urine from individuals using triclosan-containing products (8, 9). In all cases, triclosan was added 30 minutes prior to the addition of the specified bactericidal antibiotic and both antibiotics were maintained in the culture for the remainder of the experiment.

Triclosan had a dramatic protective effect on *E. coli* in an end point assay, increasing survival by several orders of magnitude in the presence of three bactericidal antibiotics and providing nearly complete protection against a fourth (Fig. 1). *E. coli* treated with triclosan exhibited a 1,000-fold increase in survival in the presence of 50 μg/mL (~5x MIC) kanamycin, an inhibitor of peptide bond formation. It also showed a 10,000-fold increase in survival in the presence of streptomycin (50 μg/mL: ~2x MIC), an inhibitor of tRNA-ribosome interaction, and ciprofloxacin (100 ng/mL: ~3x MIC) a gyrase inhibitor (Fig. 1). Strikingly, triclosan rendered *E. coli* almost completely refractory to treatment with the cell wall active antibiotic ampicillin (100 μg /mL; ~10x MIC). Viable cell numbers were essentially identical in triclosan and triclosan-ampicillin treated cultures at 2 hours, and 10% of cells in triclosan-ampicillin cultures were viable at 20 hours, suggesting triclosan increased persister frequency to all tested antibiotics.

**Figure 1:**
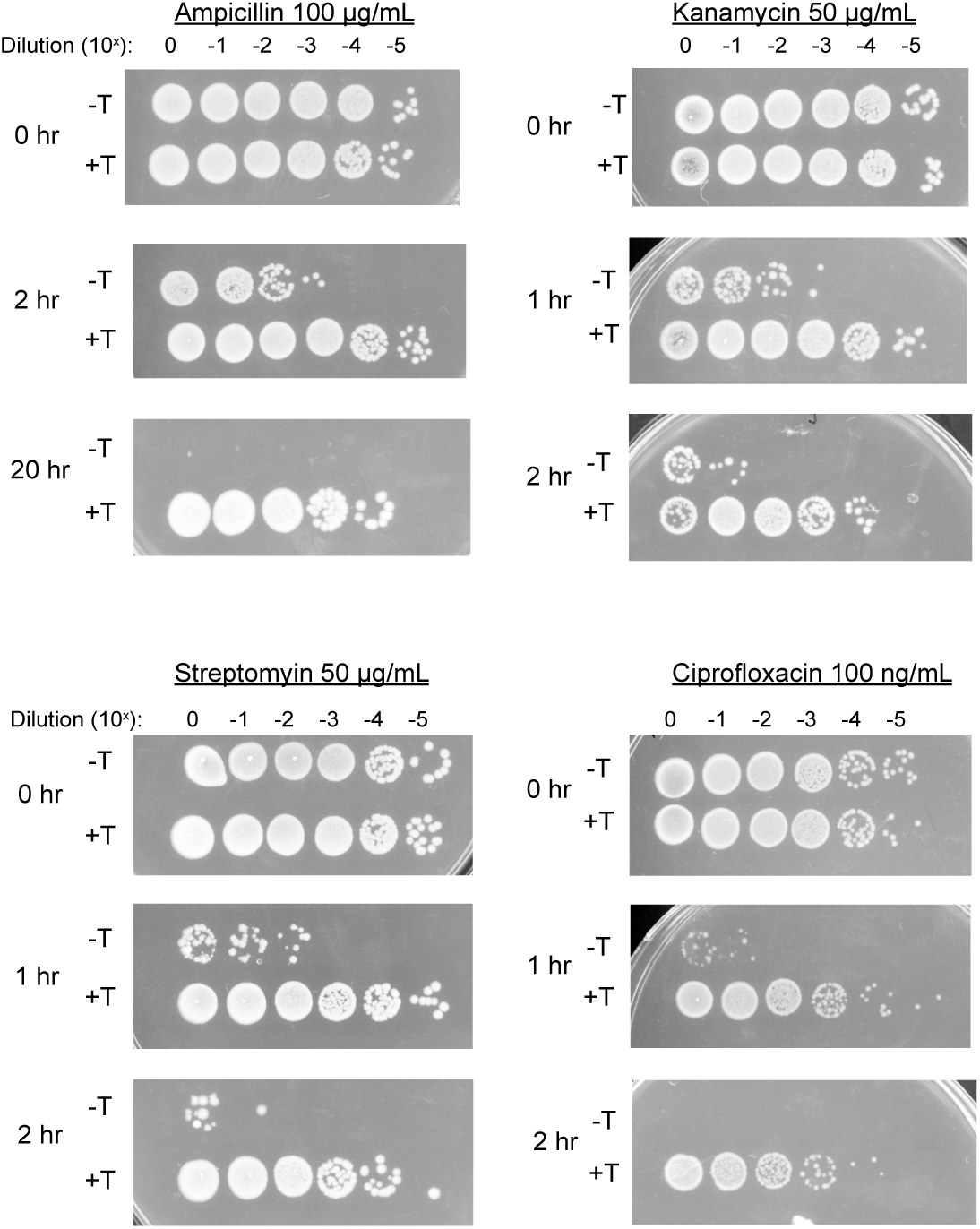
Triclosan induces tolerance to multiple antibiotics. *E. coli* (MG1655) were cultured to OD600 = 0.2, split and cultured for an additional 30 minutes with (+T) or without 200 ng/ml triclosan (-T). Indicated bactericidal antibiotics were then added and cells cultured for an additional 2 to 20 hours prior to dilution plating. Each experiment was replicated three independent times with only representative data shown.

Triclosan also protected MRSA cells from high concentrations of the glycopeptide antibiotic, vancomycin, over the course of a 20-hour experiment (Fig. 2c). MRSA treated with 100 ng/ml of triclosan were essentially refractory to 50 ng/ml vancomycin (10x the MIC) at 4 hours and exhibited a viable cell count 200x that of untreated cells at 8 hours. Even at 20 hours the viable cell count was several times higher in the presence of both triclosan and vancomycin than vancomycin alone. This delayed reduction in viable cell count is consistent with induction of a persistent state (10).

**Figure 2:**
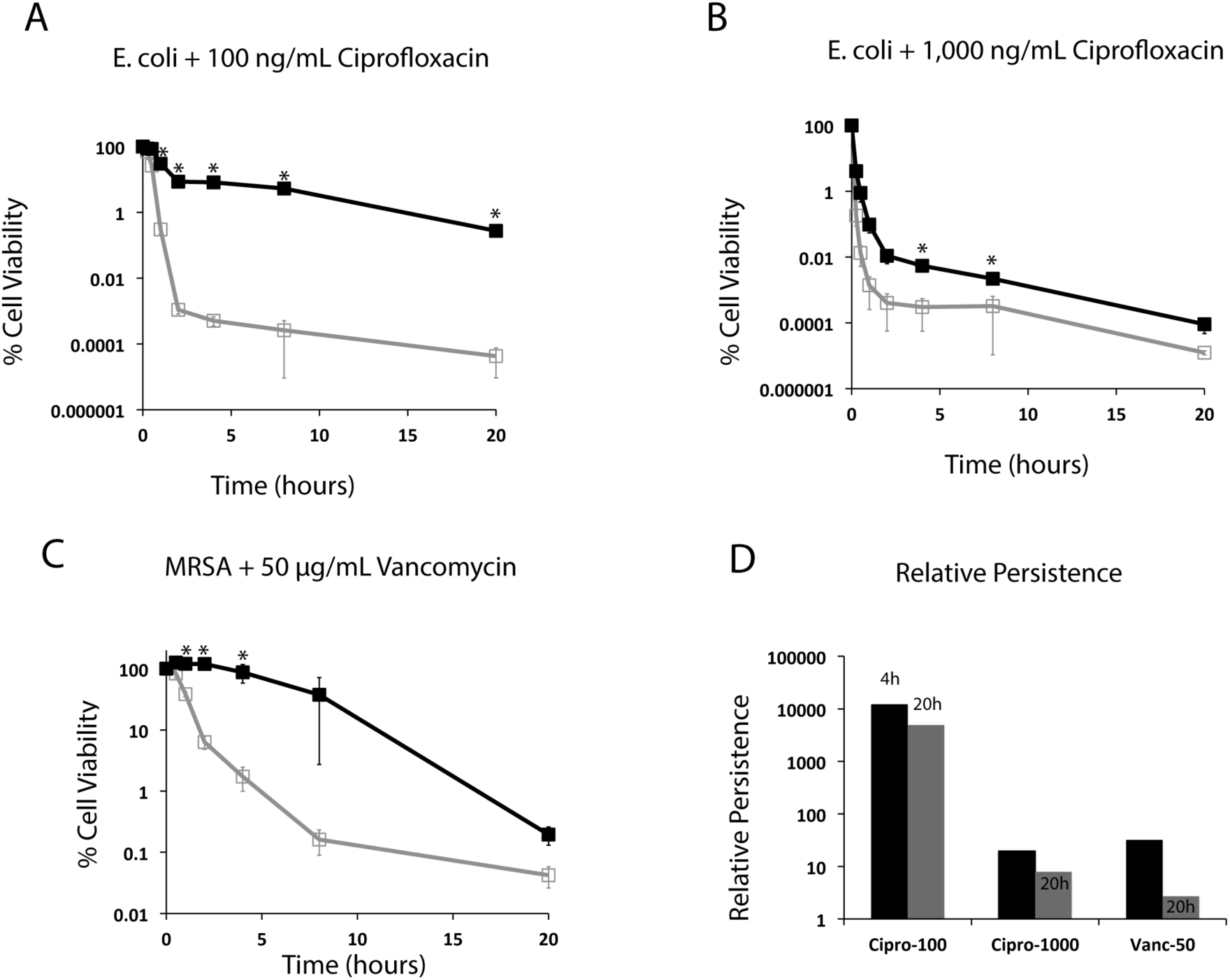
Kinetic analysis of triclosan-induced persistence. *E. coli* (MG1655) and MRSA (FPR3757) cells were cultured to OD-600 = 0.2, split and cultured for an additional 30 minutes with (black line, closed squares) or without triclosan (grey line, open squares). At t=0, 100 ng/mL (A) or 1,000 ng/mL ciprofloxacin (B) was added to *E. coli* cultures and 50 ng/ml vancomycin was added to MRSA cultures (C). Relative persistence in the presence of triclosan (CFU+T/CFU-T) was calculated from the 4- and 20-hour time points (D). Values are the mean of three independent biological replicates with error bars representing the standard error of the mean. Asterisks represent significant difference between the triclosan treated and non-treated using a Student’s two-tailed t-test with * = p< 0.05.

### Triclosan increases persister cell frequency

To further assess the protective effect of triclosan, we next performed kinetic kill curves, in which we measured colony-forming units (CFU) over a 20-hour time frame. If persister cells are present, we expect to observe two slopes, one corresponding to the kill rate of the general population and the second slope corresponding to the slower kill rate of persister cells (10). In this manner, the kill curve is able to separate antibiotic tolerance at the population level from the impact of triclosan treatment on persister levels.

For these experiments, we focused on ciprofloxacin, the broad-spectrum antibiotic used to treat urinary tract infections (UTIs). Consistent with the results of the end point assay (Fig. 1), triclosan substantially protected *E. coli* from ciprofloxacin-induced cell death throughout the duration of the time course (Fig. 2a, b). Protection was particularly pronounced at the 2-hour time point, where the slope of the kill curve for pre-treated cells diverged substantially from that of untreated cells (Fig. 2a). A reduced kill rate suggests that the pre-treated population contains a larger proportion of persister cells (21).

In agreement with previous work (16), persister population size was proportional to the concentration of ciprofloxacin. 10% of triclosan treated MG1655 cells cultured in 100 ng/mL ciprofloxacin remained viable after 2 hours (Fig. 2a), while only 0.1% cultured in the more clinically-relevant 1,000 ng/mL ciprofloxacin were viable at the same time point (Fig. 2b). For perspective, 0.1% is 1,000-fold higher than the expected frequency of persisters in an untreated population (10). After 20 hours, 90,000 CFU/mL were viable in cultures treated with both triclosan and 100 ng/mL ciprofloxacin (Fig. 2a), compared to 20 cells/mL in cultures treated with ciprofloxacin alone. At 1,000 ng/mL ciprofloxacin, cultures treated with both triclosan and ciprofloxacin contained 30 viable cells per mL (Fig. 2b). In contrast, no viable cells were detected (<10 cells/mL) in cultures treated with 1,000 ng/mL ciprofloxacin alone. We observed an increase in the abundance of persisters at drug concentrations above 1,000 ng/mL (Fig S1) (16). Although this finding initially appeared counterintuitive, it is consistent with previous reports suggesting that prophage induction in response to DNA damage is responsible for cell death at lower concentrations of ciprofloxacin, while higher concentrations kill cells before prophage are induced (16).

### Triclosan-mediated tolerance requires ppGpp

Based on the well-established connection between defects in fatty acid synthesis and accumulation of ppGpp, we speculated that triclosan-mediated tolerance was dependent on the synthesis of the alarmone (15). To test this idea, we compared the relative viability of wild-type *E. coli* and mutants unable to synthesize the alarmone (ppGpp0; *spoT::cat* Δ*relA*) 2 hours after antibiotic challenge in the presence or absence of triclosan.

Although triclosan inhibited the growth of both wild-type and ppGpp0 cells, it was unable to substantially protect ppGpp0 cells from any of the four bactericidal antibiotics we tested. These include ampicillin and ciprofloxacin, as well as the translation inhibitors, kanamycin and streptomycin (Fig. 3a, b). The ppGpp0 cells were more sensitive to kanamycin and streptomycin, showing no viable cells after 60-minute treatment, thus measurements were performed at 30 minutes. Importantly, triclosan alone was not bactericidal to either wild-type or ppGpp0 cells (Fig. S2).

In contrast to triclosan, pre-treatment with another bacteriostatic compound, spectinomycin, increased tolerance to kanamycin, streptomycin, ampicillin, and ciprofloxacin in both wild-type and ppGpp0 mutant cells (Fig. 3C). A translation inhibitor, spectinomycin does not impact ppGpp levels in *E. coli* (20). While spectinomycin was still protective in the ppGpp0 cells, levels of protection were slightly decreased compared to WT cells.

**Figure 3.**
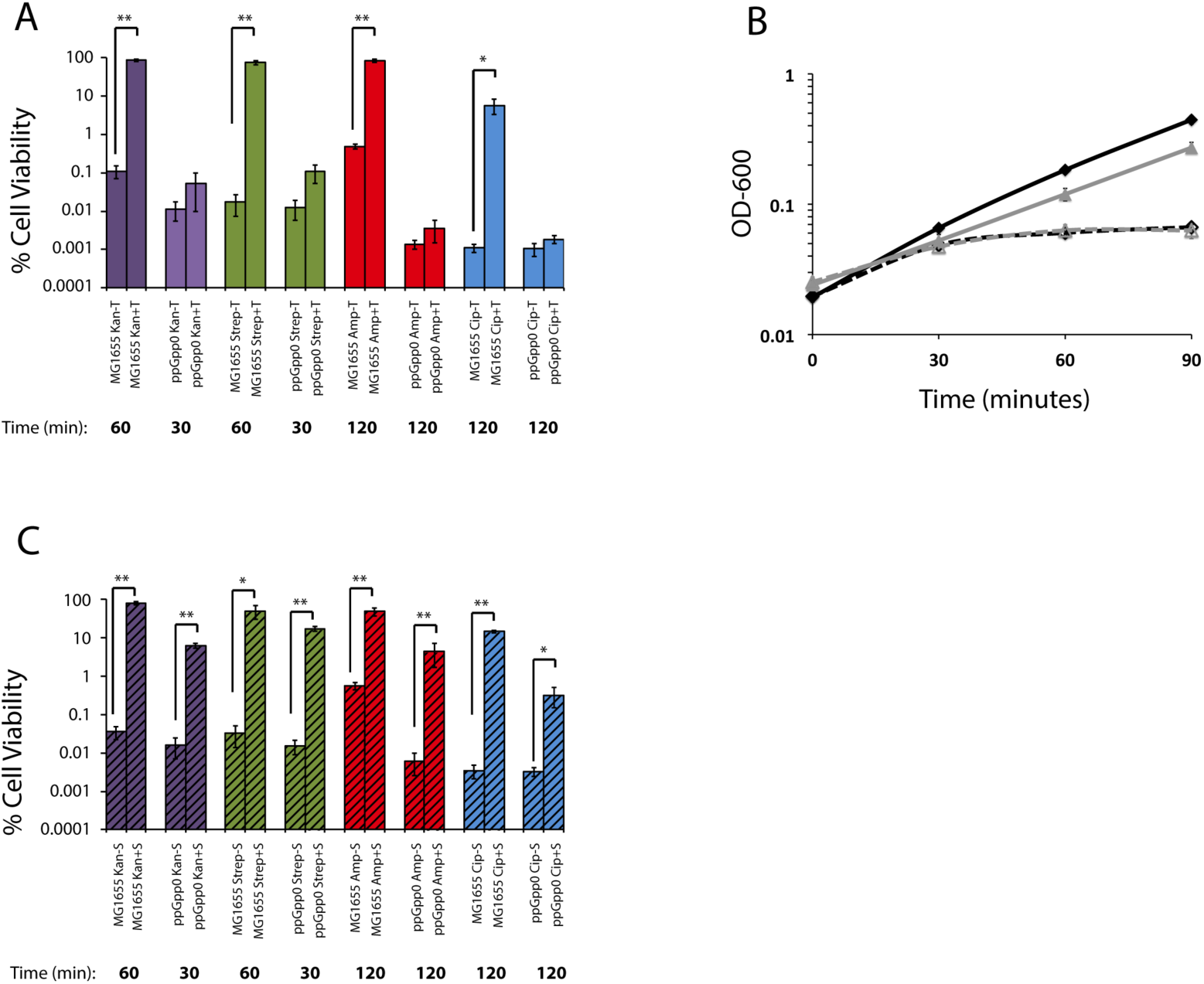
ppGpp is needed for triclosan induced tolerance. Cell viability of MG1655 and ppGpp0 *E. coli* with (+T) or without (-T) pretreatment with triclosan after challenge with antibiotic (A). Growth curves of MG1655 (black curve) or ppGpp^0^ (gray curve) in LB with (dashed lines) or without (solid lines) triclosan (B). Cell viability of MG1655 and ppGpp0 *E. coli* with (+T) or without (-T) pretreatment with spectinomycin after challenge with antibiotic (C). Values are the mean of three independent biological replicates with error bars representing the standard error of the mean. Asterisks represent significant difference between the triclosan treated and non-treated using a Student’s two-tailed t-test with * = p< 0.05 and ** = p<.001.

### Triclosan drives tolerance to ciprofloxacin in a murine model

Due to its widespread use and inherent stability, triclosan is present in both human populations and the environment (8). Thus, a key question is whether the tolerance we observed *in vitro* is relevant *in vivo*. To determine the physiological relevance of triclosan-mediated tolerance, we employed a mouse model of *E. coli* UTI. UTIs are one of the most prevalent bacterial infections, impacting approximately 150 million people annually (22). Uropathogenic *E. coli* (UPEC) is the main causative agent of both uncomplicated and complicated UTI (23). Pretreatment with triclosan rendered the well-characterized *E. coli* cystitis isolate UTI89 ~10-fold more tolerant to 1,000 ng/mL ciprofloxacin than untreated cells at 2 hours, a level equivalent to the tolerance we observed for *E. coli* MG1655 at the same time point (Fig. 1 and Fig. S3).

For *in vivo* experiments, we provided six-week old female wild-type C3H/HeN mice with drinking water containing 1,000 ng/mL triclosan for 21 days. Control mice were given plain water for the same duration. At 21 days, experimental and control mice were trans-urethrally infected with ~5 × 10^7^ CFU of *E. coli* UTI89. At 24 hours post-infection, a subset of the mice was treated with intraperitoneal ciprofloxacin (25 mg/kg). At 48 hours post-infection, all mice were sacrificed and bacterial colonization assessed in urine and bladder.

After ciprofloxacin treatment, bacterial titers were >100-fold higher in the urine (*p*<0.0001) and >10-fold higher (*p*<0.0001) in the bladders of triclosan-treated mice versus control animals (Fig. 4a, b), consistent with triclosan-induced tolerance occurring *in vivo*. Bacterial load at 24 hours post-infection was nearly equivalent in triclosan-treated and control mice, indicating that triclosan did not significantly impair UTI89 viability (Fig. 4a, b). Treated mice had triclosan levels between 70–750 ng/mL in their urine, comparable to the MIC for *E. coli* (200 ng/mL) and similar to reported triclosan levels in human urine (2.4 to 3,790 ng/mL) (8) (Fig. 4c). We also detected two putative metabolized forms of triclosan, one with a mass consistent with the previously reported sulfonated triclosan (24), and the other 96 daltons larger (Fig. S4). Whether or not the modified forms of the drug are active against bacteria is unclear. Control mice had no observable triclosan (below 1.6 ng/mL limit of detection).

**Figure 4:**
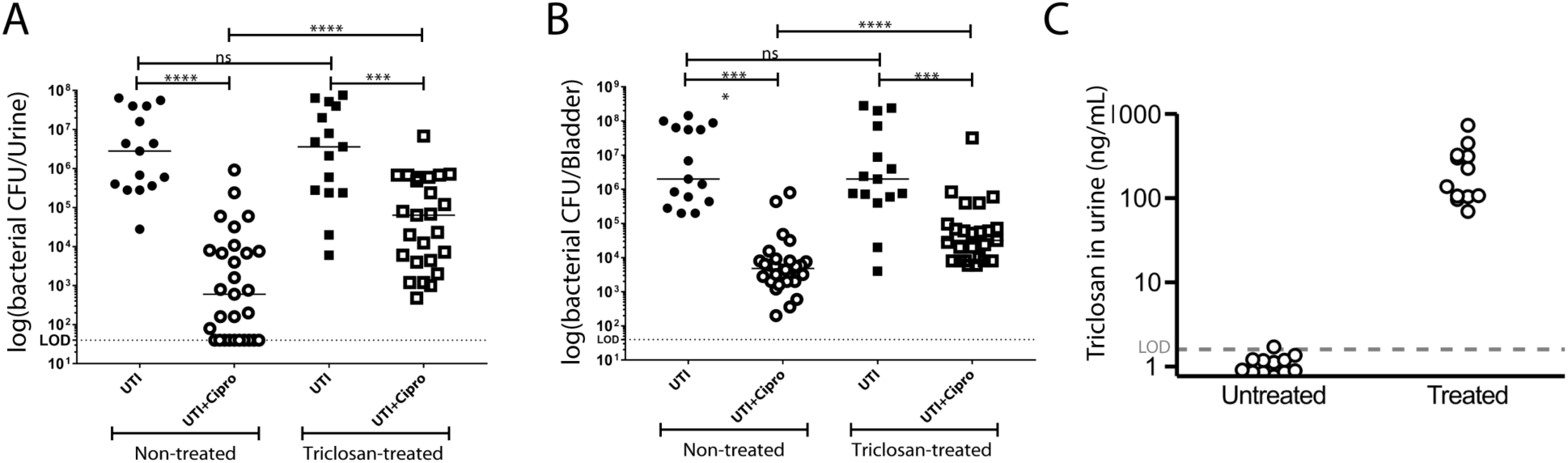
Triclosan reduces ciprofloxacin efficacy up to 100-fold in a mouse UTI model. For each round of the experiment, 15 mice were given water containing triclosan (1,000 ng/mL) for 21 days, and 15 control mice received plain water. At 21 days, all mice were infected with *E. coli* UTI89 (~5 × 10^7^ CFU). 24 hours post-infection, 10 mice from each group were given 25 mg/kg of ciprofloxacin intraperitoneally. 48 hours post infection, urine (A) was taken and mice were sacrificed and bladders (B) were harvested. To compare the mice groups, Mann-Whitney U test was used, p<0.05 was considered statistically significant: *, p<0.05; **, p<0.005; ***, p<0.0005; ns, values were not statistically different. The horizontal bar represents the median value. The horizontal broken line represents the limit of detection (LOD) of viable bacteria. Data are from 3 independent experiments. (C) Free triclosan levels measured by LC-MS/MS of triclosan in untreated and treated mouse urine. (45 mice were untreated and 45 mice were treated, these mice were divided in three independent experiments. Urine from 3–4 mice was pooled for triclosan analysis). The horizontal broken line represents the limit of detection (LOD) of triclosan.

## Discussion

Our data indicate that environmentally relevant concentrations of triclosan reduce antibiotic efficacy as much a 100-fold *in vivo* (Fig. 4) and highlight an unexpected and potentially important role for triclosan as a contributor to antibiotic tolerance and bacterial persistence in both community and healthcare settings. Triclosan-mediated tolerance is dependent on ppGpp synthesis, most likely in response to inhibition of fatty acid synthesis (Fig. 3a) (15). This finding is consistent with prior work implicating ppGpp in antibiotic tolerance and persister development (16).

In contrast to previous studies of ppGpp-induced persistence that relied on either carbon starvation or the addition of serine hydroxamate to induce accumulation of high concentrations of ppGpp (100x above baseline) (25, 26), defects in fatty acid synthesis have at best a modest impact on ppGpp levels (~5x over baseline) (15). This suggests that even relatively low levels of ppGpp are sufficient to protect cells from a panel of antimicrobials. Specifically how modest increases in ppGpp might confer tolerance to different antibiotics thus remains an open question.

We favor the hypothesis that ppGpp mediates changes in individual biosynthetic pathways that render them tolerant of their cognate antimicrobial. For example, ppGpp-dependent down-regulation of ribosomal RNA synthesis significantly curtails translation (27), potentially conferring tolerance to the translational inhibitors kanamycin and streptomycin. Similarly, increases in ppGpp are reported to curtail DNA replication (28)—both elongation and initiation—providing a straightforward explanation for ppGpp-mediated ciprofloxacin resistance.

It is generally recognized as poor practice to prescribe a bacteriostatic compound prior to or along with delivery of a bactericidal one (29), because of the potential that the former will interfere with the activity of the latter. At the same time, their mechanisms of action, and therefore the mechanisms by which these drugs drive tolerance, are likely to differ widely. While the translation inhibitor spectinomycin provides protection against bactericidal compounds, it does not induce accumulation of ppGpp (20), a fact supported by our finding that spectinomycin induces tolerance to bactericidal compounds in both wild-type and ppGpp0 cells (Fig. 3c). At the same time, ppGpp-dependent induction of antibiotic tolerance is likely to be a feature triclosan shares with a related compound, triclocarban, which also inhibits an early step in fatty acid synthesis, and is also a common additive in consumer products. Triclosan also stands out from other bacteriostatic compounds by virtue of its widespread use and sheer abundance in the environment. ~1 kg of Triclosan is produced for every 3 kg of other antimicrobials and estimates indicate that ~100 metric tonnes are being deposited annually in the environment through waste water treatment in the US alone (30).

Although triclosan has low toxicity (LD-50 4,350 mg/kg orally) (31), accumulating data link long-term exposure with antibiotic resistance (32) and there are reports that triclosan may also function as an endocrine disrupter (33, 34). Our analysis of the impact of triclosan on antibiotic efficacy in a mouse UTI model (Fig. 4) highlights yet another deleterious “side effect” of this ubiquitous antimicrobial. UTIs alone impact 150 million people worldwide (22) at a cost of $3.5 billion per year in the US alone (35). Complications associated with UTIs include pyelonephritis with sepsis, renal damage, pre-term birth, *Clostridium difficile* colitis, sepsis, and death particularly in the very old and the very young (23). Coupled with the well established connection between antibiotic tolerance and recurrent/chronic infections (12, 13), our findings reinforce the need for substantial caution—as well as consideration of unintended consequences—in evaluating the costs and benefits of antimicrobial additives in consumer products.

## Methods

### Materials and Strains

Triclosan, ampicillin, kanamycin, streptomycin, ciprofloxacin, and vancomycin were purchased from Sigma-Aldrich. Stock solutions were made in water for ampicillin (100 mg/mL), kanamycin (50 mg/mL), streptomycin (100 mg/mL) and ciprofloxacin (10 mg/mL). Triclosan was dissolved in ethanol (10 mg/mL), and vancomycin was dissolved in DMSO (100 mg/mL). *E. coli* MG1655 and *S. aureus* FPR3575 were both lab strains and *E. coli* UTI89 was isolated from a patient with a urinary tract infection (36). *E. coli* were grown in Luria-Bertani broth (LB) and *S. aureus* was grown in tryptic soy broth (TSB). Growth temperature was 37 °C for all experiments.

### Determination of Minimum Inhibitory Concentration (MIC)

To determine the MIC for the panel of antibiotics utilized in this study, *E. coli* and *S. aureus* were grown to OD-600 = 0.1 in LB or TSB, respectively. Cells were then back-diluted 1,000-fold and transferred to a 96-well plate containing 2-fold dilutions of respective antibiotics and cultured at 37 °C for 16 additional hours with vigorous shaking in a BioTek Eon plate reader. MIC was calculated as the lowest antibiotic concentration, preventing development of detectable turbidity at OD-600.

### Assays for antibiotic tolerance and persistence

To assay tolerance and persistence, *E. coli* and *S. aureus* were grown to an OD-600 = 0.2 in LB or TSB, respectively. Cells were then back-diluted to an OD-600 = 0.1 in media containing triclosan at indicated concentrations and cultured for an additional 30 minutes, before being challenged with bactericidal antibiotics. For dot plating, 10 μL of a 10-fold dilution series was plated on antibiotic-free LB agar or TSB agar as appropriate. For determination of colony forming units (CFU), 100 μL of a 10-fold dilution series was spread on antibiotic-free LB agar or TSB agar plates. Cells were incubated for ~12 hours at 37 °C prior to quantification. CFUs were normalized to CFUs at the initial time point to correct for the ~2-fold increase in cell number in untreated cultures during the 30-minute pre-treatment period. Relative persistence is defined as the CFU’s of the triclosan treated sample divided by the CFU’s of the non-treated sample.

### UTI mouse work

Six-week old female wild-type C3H/HeN mice were obtained from Envigo. Mice were treated with or without 1,000 ng/mL (100 ppm) triclosan in the drinking water for 21 days. At 21 days, mice were anesthetized by inhalation of 4% isoflurane and mouse bladders were trans-urethrally infected with approximately 5 × 10^7^ CFU of *E. coli* UTI89 in 50μl PBS(37). Briefly, a single UTI89 colony was inoculated in 20 ml of Luria Broth (LB) and incubated at 37 °C under static conditions for 24 hours. Bacteria were then diluted (1:1,000) into fresh LB and incubated at 37 °C under static conditions for 18 to 24 hours. Bacteria were subsequently washed three times with PBS and then concentrated to approximately 5 × 10^7^ CFU per 50 μL. At 24 hours post-infection, mice received 25mg/kg of ciprofloxacin intraperitoneal. 48 hours post-infection, mice were euthanized, and bladders were harvested and urine was collected. Bladders were homogenized in PBS and bacterial load present in bladders and urines was determined by plating serial dilutions on LB agar supplemented with antibiotics when appropriate. Statistical analyses were performed using the Mann–Whitney U test with GraphPad Prism software (version 6.0 for Mac). All animal studies were performed in accordance with the guidelines of the Committee for Animal Studies at Washington University School of Medicine.

### Measurement of triclosan and metabolites in mouse urine

Since triclosan has been observed to adsorb to plastic surfaces, sample handling was performed in glass vessels whenever possible (24). A stock solution of 1 mg/mL triclosan (Sigma) was prepared in methanol and a 100 μg/mL ^13^C_12_-triclosan (99%) internal standard in MTBE was purchased from Cambridge Isotope Laboratories (Andover, MA). A dilution series of 1,000, 200, 40, 8, 1.6, and 0.32 ng/mL triclosan was prepared in pooled, untreated mouse urine and spiked with 100 ng/mL ^13^C_12_-triclosan internal standard. Samples were diluted 1:1 in methanol, spun down at 20,000×g for 10 minutes, and filtered through 0.45 μm 13 mm diameter PVDF syringe filters (Millipore). Finally, cleaned samples were diluted 1:1 in HPLC-grade water (Sigma).

Using a Shimadzu UFLC (Kyoto, Japan), 10 μL of each sample was injected onto a fused core phenyl-hexyl column (100 mm × 2 mm × 2.7 μm) with a 0.4 mL/min flow rate (Ascentis Express, Supelco). Triclosan was eluted from the column as follows: Solvent A (0.1% formic acid) and Solvent B (90% acetonitrile with 0.1% formic acid) were held constant at 80% and 20%, respectively, for 0.1 minutes. Solvent B was increased to 98% by 5 minutes, held at 98% for 1 minutes, and then reduced again to 20% in 0.1 minutes. The column was equilibrated in 20% Solvent B for 3 minutes between runs.

Triclosan was detected using an AB Sciex API 4000 QTrap mass spectrometer (AB Sciex, Foster City, CA) running in negative ion electrospray ionization mode (ESI) using a Turbo V ESI ion source. Triclosan was detected using the instrument settings listed in Supplementary Data Table 1. A precursor ion scan was performed for the 35 m/z product ion to determine the mass spectrum of triclosan, ^13^C_12_-triclosan, and any potential metabolites (Fig. S4a). Because triclosan contains three chlorine atoms, its mass spectrum includes prominent isotope peaks (M+2, M+4) corresponding to the natural abundance of ^37^Cl (Fig. S4b). To improve sensitivity, product ions from the two most abundant isotopologues were detected and added together prior to peak integration. Peaks for triclosan and internal standard were integrated with Analyst software (AB Sciex) and normalized. Normalized peak areas varied linearly with triclosan concentration above 1.6 ng/mL.

Pooled urines from 3 to 4 mice were spiked with 100 ng/mL internal standard and cleaned as described above. Samples were analyzed by LC-MS/MS and triclosan was quantified using the standard curve (Fig. S4e).

### Statistical analysis

Values for the *in vitro* data are expressed as the mean ± standard error of the mean from n=3 replicates. *In vitro* data was analyzed using a two-tailed Student’s t-test with statistical significance determined when p<.05. For the mouse data, the Mann-Whitney U test was used to test for statistical significance. Values represent means ± SEM derived from at least 3 independent experiments. *, *P*<0.05; **, *P*<0.005; ***, *P*<0.0005; ****, *P*<0.00005; ns, difference not significant.

## Acknowledgements

This work was supported by National Institutes of Health grants R01-GM64671 and R35-GM127331 to P.A.L. and R01-DK051406, R01-AI108749-01, and P50-DK0645400 grants to A.L.F-M and S.J.H. C.S.W was supported in part by an Arnold O. Beckman Postdoctoral Fellowship. J.I.R. and J.P.H. were supported by National Institutes of Health grant RO1DK111930. The authors thank Dr. Robyn Klein and members of the Levin, Hultgren, and Zaher groups for insightful suggestions and critical comments on the manuscript.

## Author Contributions

CSW, PAL, ALFM, SJH, JIR and JPH designed the research studies. CSW performed the *in vitro* assays. ALFM and AJLL performed the animal experiments and acquired data. JIR and JPH performed triclosan detection and quantification experiments and acquired data. CSW ALFM, JIR, and PAL analyzed data. CSW, ALFM, and JIR prepared the figures. CSW, PAL, and ALFM wrote the manuscript. CSW, PAL, ALFM, SJH, JIR and JPH reviewed and edited the manuscript.

## Supplementary Data

**Supplementary Data Figure 1.**
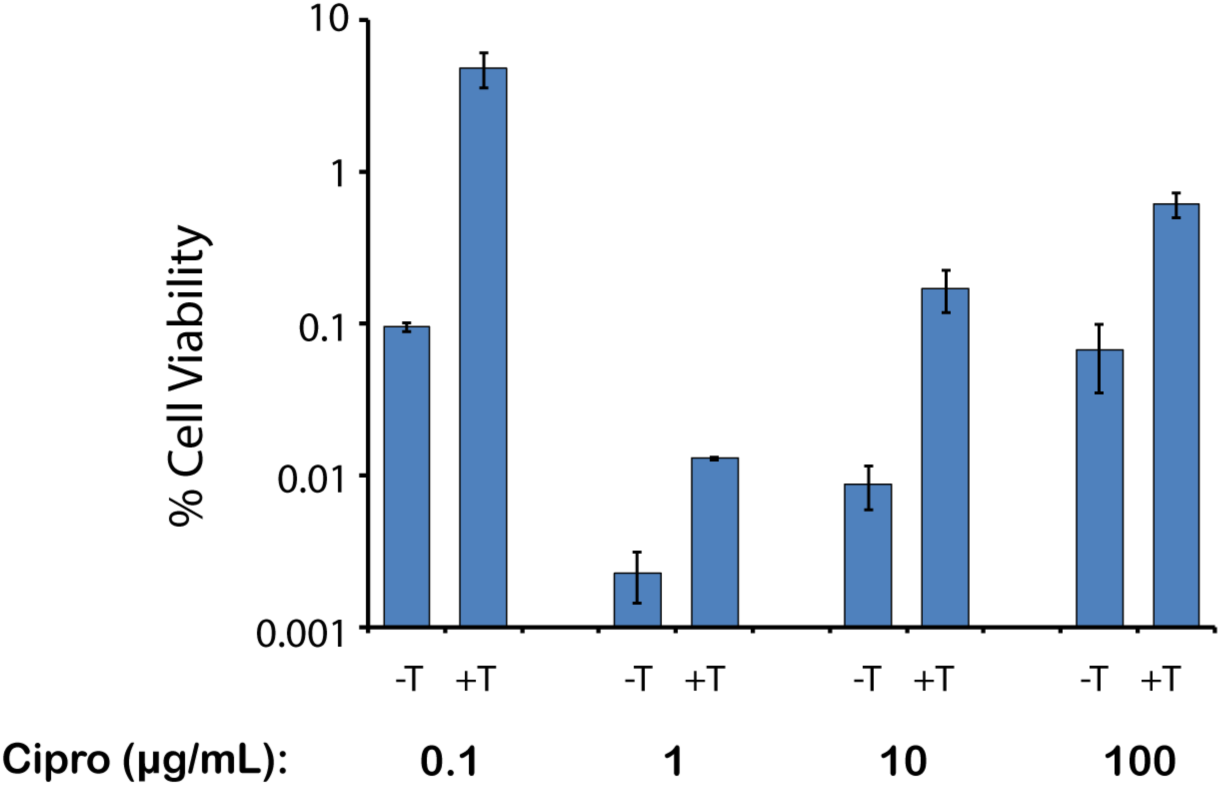
Triclosan protects at high concentrations of ciprofloxacin. MG1655 cells were grown to OD-600 = 0.1 before triclosan was added for a final concentration of 200 ng/mL for 30 minutes. Ciprofloxacin was added at the labeled concentration. Ciprofloxacin was washed off, and cell viability was determined. Values are shown as averages of three replicates with error bars showing the standard error of the mean.

**Supplementary Data Figure 2.**
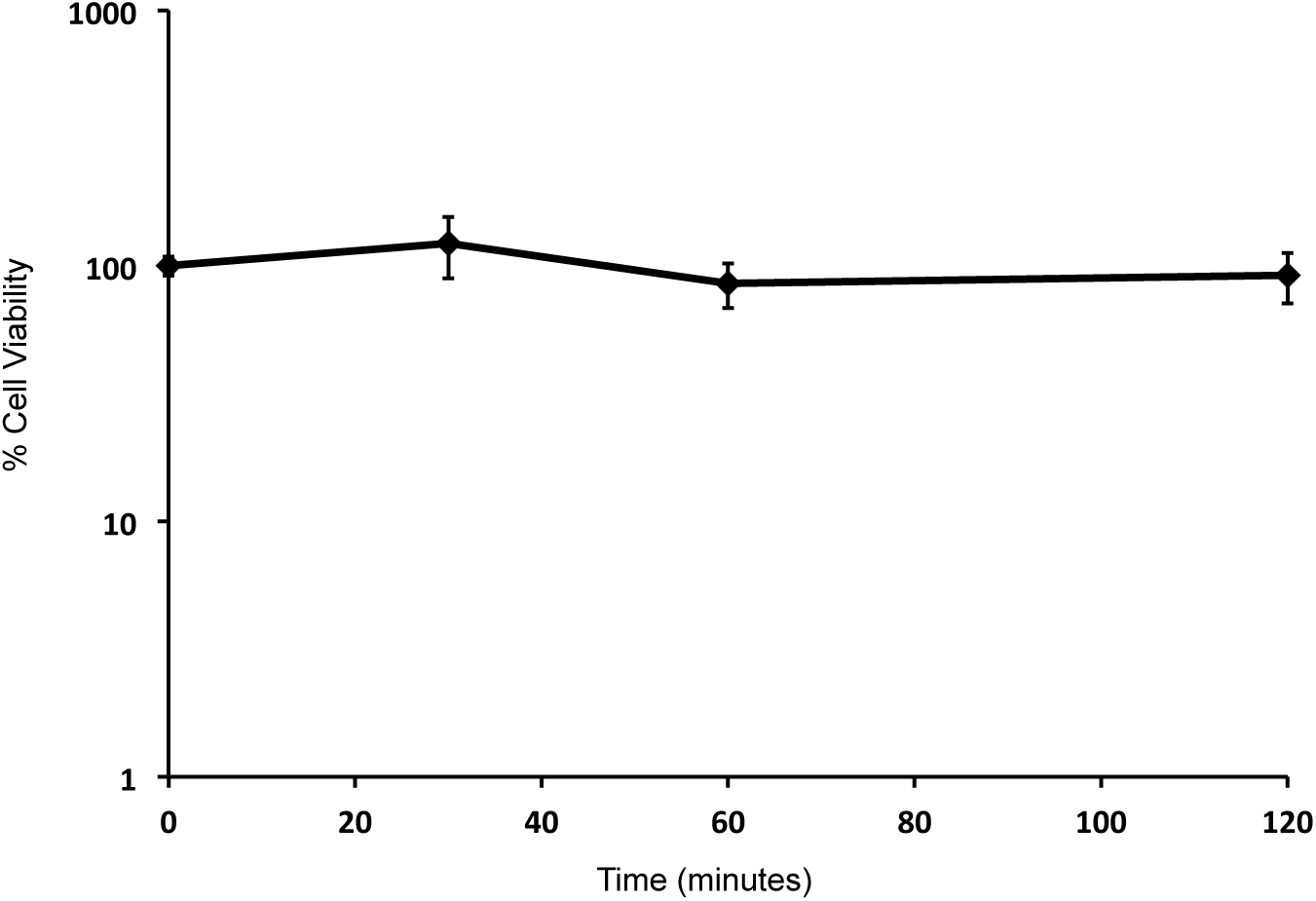
Triclosan is not bactericidal to the ppGpp0 cells. ppGpp0 cells were grown to OD-600 = 0.2 before triclosan was added for a final concentration of 200 ng/mL. Cells were plated at each time pointed and colony-forming units were quantified. Each point represents the average of three biological replicates with the error bars representing the standard error of the mean.

**Supplementary Data Figure 3.**
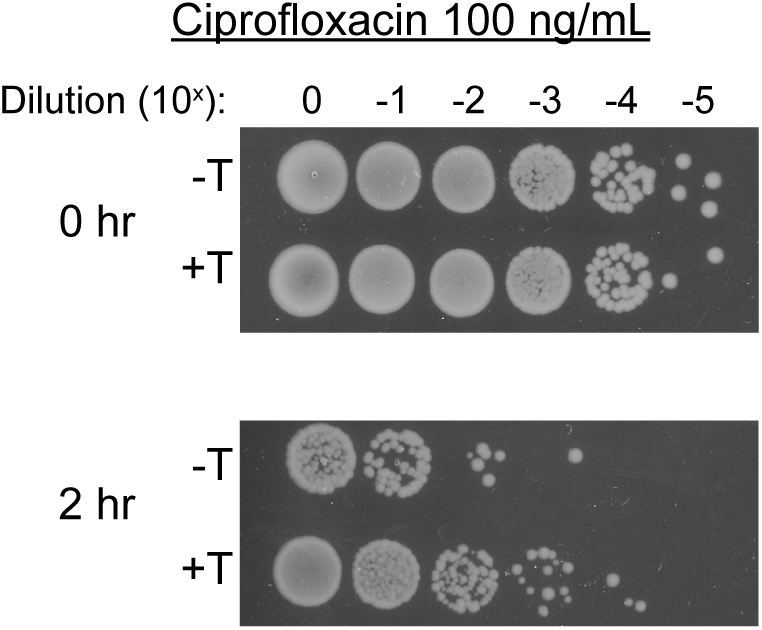
Triclosan induces ciprofloxacin tolerance in the uropathogentic *E. coli* UTI89. UTI89 cells were grown up to OD-600 = 0.2, split and cultured for an additional 30 minutes with (+T) or without 200ng/ml triclosan (-T). Ciprofloxacin was added to obtain a final concentration of 100 ng/mL. Cells were dot-plated at the 0- and 2-hour time points. The plating efficiency was repeated three independent times with a representative image shown.

**Supplementary Data Figure 4.**
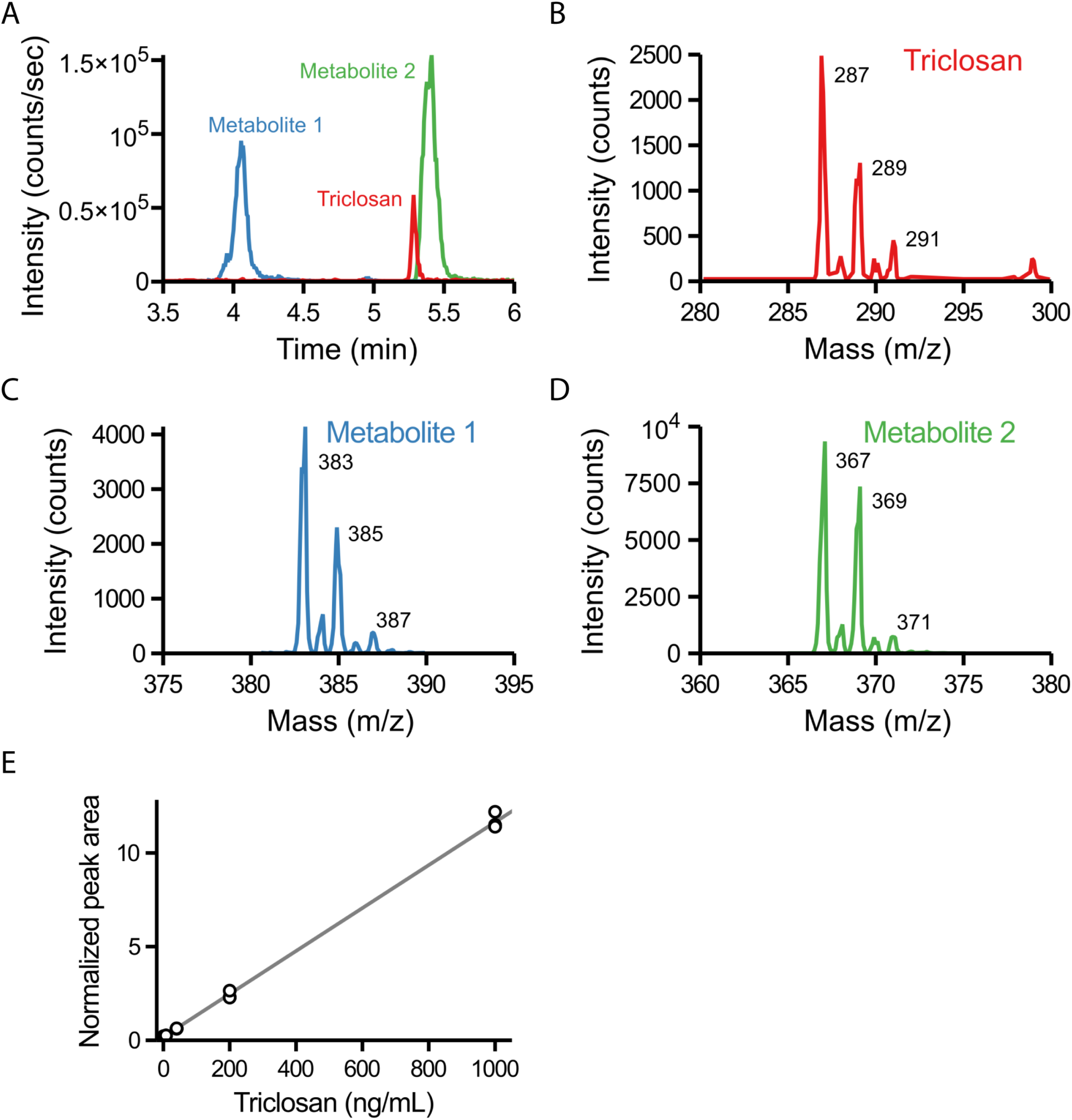
Measurement of free and metabolized triclosan in triclosan-treated mouse urine. Chromatogram of a representative LC-MS/MS experiment showing the elution profile of metabolite 1 (blue), triclosan (red) and metabolite 2 (green) (A). Mass spectra of triclosan, metabolite 1, and metabolite 2, respectively, measured using precursor ion scans for the chloride product ion (35 m/z) (B, C, and D). Triclosan peak area detected by LC-MS/MS varies linearly with concentration down to 1.6 ng/mL (E).

**Supplementary Data Table 1.** MS/MS settings for triclosan detection.

| Instrument Settings |  |  |
| --- | --- | --- |
| Ion Spray Voltage | -4.5 kV |  |
| Heater Temperature | 500 °C |  |
| Nebulizer Gas | 40 |  |
| Auxiliary Gas | 40 |  |
| Declustering Potential | -10 V |  |
| Collision Energy | -40 V |  |
| Ions Detected (m/z) |  |  |
| Triclosan | 286.8 → 35 | 288.7 → 35 |
| <sup>13</sup>C<sub>12</sub>-Triclosan | 298.84 → 35 | 300.74 → 35 |
| Metabolite 1 | 382.8 → 35 | 384.8 → 35 |
| Metabolite 2 | 366.7 → 35 | 368.8 → 35 |

